# DNA replication initiation timing is important for maintaining genome integrity

**DOI:** 10.1101/2024.06.18.599555

**Authors:** Tristan T. Reed, Abigail H. Kendal, Katherine J. Wozniak, Lyle A. Simmons

## Abstract

DNA replication is regulated by factors that promote or inhibit initiation. In *Bacillus subtilis,* YabA is a negative regulator of DNA replication initiation while the newly identified kinase CcrZ is a positive regulator. The consequences of under-initiation or over-initiation of DNA replication to genome stability remain unclear. In this work, we measure origin to terminus ratios as a proxy for replication initiation activity. We show that Δ*ccrZ* and several *ccrZ* alleles under-initiate DNA replication while ablation of *yabA* or overproduction of CcrZ leads to over-initiation. We find that cells under-initiating DNA replication have few incidents of replication fork stress as determined by low formation of RecA-GFP foci compared with wild type. In contrast, cells over-initiating DNA replication show levels of RecA-GFP foci formation analogous to cells directly challenged with DNA damaging agents. We show that cells under-initiating and over-initiating DNA replication were both sensitive to mitomycin C and that changes in replication initiation frequency cause increased sensitivity to genotoxic stress. With these results, we propose that cells under-initiating DNA replication are sensitive to DNA damage due to a shortage of DNA for repair through homologous recombination. For cells over-initiating DNA replication, we propose that an increase in the number of replication forks leads to replication fork stress which is further exacerbated by chromosomal DNA damage. Together, our study shows that DNA replication initiation frequency must be tightly controlled as changes in initiation influence replication fork fate and the capacity of cells to efficiently repair damage to their genetic material.

**IMPORTANCE:** The regulation of DNA replication is fundamental to cell growth and cell cycle control. In eukaryotes under-initiation or over-initiation leads to genome instability. For bacteria, it is unclear how changes in replication initiation frequency impact DNA replication status and genome integrity. We show that tight regulation of DNA replication initiation is critical for maintaining genome integrity. Cells over-initiating or under-initiating DNA replication are sensitive to DNA damage. Further, cells over-initiating DNA replication experience replication fork stress at levels that phenocopy cells encountering DNA damage from the crosslinking agent mitomycin C. Our results establish the critical importance of properly regulating DNA replication initiation frequency because an imbalance in initiation results in replication fork perturbations, deficiencies in DNA repair, and genome instability.

## INTRODUCTION

All organisms are required to replicate their genomic DNA at the proper time during growth or the appropriate stage of the cell cycle. To ensure that replication occurs at the proper time, DNA replication initiation is a carefully regulated process. DNA replication initiation in bacteria and eukaryotes is mediated by initiation proteins that are part of a large superfamily of <u>A</u>TPases <u>a</u>ssociated with diverse cellular <u>a</u>ctivities (AAA+) (1–5). In bacteria, DnaA is a highly conserved AAA+ superfamily protein required for replication initiation from the origin, *oriC* (3, 6). DnaA binds ATP and then DnaA boxes located within *oriC* (3, 6). Once bound, DnaA unwinds the AT-rich region of *oriC* and associates with helicase loader proteins (2, 7). After the helicase is loaded, the remaining proteins assemble to allow for priming and bidirectional replication of the chromosome (2, 8).

Because DnaA represents the committed step to replication initiation, DnaA is subject to positive and negative regulation (3). In *E. coli* DnaA is negatively regulated by processes that limit its ATP binding or hydrolysis (9–11), while in the Gram-positive bacterium *B. subtilis,* negative regulators tend to disrupt DnaA filament formation *oriC* (12–14). YabA and Soj/ParA are examples of negative regulators of DNA replication in *B. subtilis* that interfere with the ability of DnaA to form a nucleoprotein filament at the *oriC* (12, 13, 15–18). YabA has been shown to bind DnaA and the replication sliding clamp DnaN (14, 18). Binding of YabA to DnaA interferes with the cooperative binding of DnaA to *oriC,* inhibiting initiation (14, 18). Therefore, cells bearing a *yabA* null allele over-initiate DNA replication because negative control of DnaA is relieved in the absence of YabA (14–16).

In *E. coli,* there are multiple examples of DnaA variants that cause hyperactive initiation of DNA replication (11, 19, 20). The most well studied example is DnaAcos, which confers a cold sensitive phenotype caused by over-initiation of DNA replication (20–23). Studies of over-initiation of DNA replication in *E. coli* have shown that over-initiation can lead to enrichment of replication forks in origin proximal DNA, suggesting that forks have slowed, stalled, or become blocked following excessive initiation events (19). In *B. subtilis,* the *dnaA1* allele (*S401F*) is defective for replication initiation at the nonpermissive temperature (16, 24). Further, the *dnaA1* allele was recently shown to cause over-initiation of DNA replication at the permissive temperature (16). Combining *dnaA1* and Δ*yabA* resulted in a synthetic lethal phenotype, demonstrating that excessive over-initiation in *B. subtilis* results in cell death (16). In eukaryotes, over-initiation is caused by overexpression of oncogenes, cyclin E and Ras, resulting in genome instability through chromosomal breakage at fragile sites, and collisions between replication forks and transcription complexes (25–27). Furthermore, in eukaryotes under-replication also leads to genome instability including DNA breakage at fragile sites (28–30). In the case of eukaryotes, DNA replication initiation occurs once per cell cycle. For bacteria, including *E. coli* and *B. subtilis,* multiple initiation events occur during each cell cycle when cells are grown in rich medium, allowing for an increase in growth rate as new cells inherit chromosomes with forks having already initiated at the origin before cell division occurs [(31) for review (32, 33)]. The effects of under-initiation and over-initiation on genome integrity in bacterial cells using multi-fork replication remain unclear.

In *E. coli* there are a number of proteins that can help stimulate initiation activity of DnaA, including IHF and Fis [for review(3)]. CcrZ is a recently identified kinase in *B. subtilis* that stimulates DNA replication initiation through *oriC* (34). CcrZ is conserved in the phylum Firmicutes and has been shown to be important for positively regulating initiation in *Streptococcus pneumoniae*, *Staphylococcus aureus,* and *B. subtilis* (34). In each of the organisms tested, deletion or depletion of *ccrZ* resulted in a lower *oriC/ter* ratio than measured in wild type (WT) cells (16, 34). Integrity of the kinase active site of CcrZ is required for function, and CcrZ has been shown to bind DnaA and the *B. subtilis* helicase loader protein DnaB (34, 35). The cognate substrate of CcrZ remains unknown; however, CcrZ was shown to generate ADP from ATP on ribose and 2-deoxyribose, suggesting the CcrZ phosphorylates a small signaling molecule (35). Ectopic expression of *ccrZ* from a xylose inducible promoter has been shown to result in hyperactive initiation of DNA replication, sensitivity to DNA damage, and induction of the SOS response in a subpopulation of cells (35). Unlike the DnaAcos variant from *E. coli,* which leads to clear replication fork congestion near the origin, ectopic expression of CcrZ in *B. subtilis* increased initiation frequency but did not lead to replication fork pile up near the origin or anywhere else on the chromosome (19, 35). We took advantage of the multiple methods to cause over or under-initiation of replication in *B. subtilis* to determine how changes in initiation relate to genome instability.

We show that replication fork integrity is correlated with the frequency of replication initiation: decreased initiation leads to fewer fork problems while increased initiation results in fork stress consistent with cells incurring significant chromosomal DNA damage. We also show that under-initiation results in a failure to efficiently repair DNA damage, resulting in sensitivity to genotoxic stress. With our data, we suggest that under-initiation leads to DNA damage sensitivity as cells lack sufficient chromosomal DNA to efficiently engage in homologous recombination. Together, our work shows that initiation frequency is carefully regulated to safeguard the genome from replication fork problems and to allow for efficient DNA repair when genome integrity is compromised.

## RESULTS

### Overproduction of CcrZ bypasses negative regulation and leads to excessive over-initiation of DNA replication

The rationale for this work is to understand the effect of over and under-initiation of DNA replication on genome integrity. Prior work from our lab showed that overexpression of *ccrZ* led to over-initiation of DNA replication as determined by whole genome resequencing (35). Work by other groups has shown that deletion of *ccrZ* results in under-initiation of DNA replication in *B. subtilis*, *S. pneumoniae,* and *S. aureus,* suggesting that the contribution of CcrZ to DNA replication initiation is conserved (16, 34). To begin, we used the same growth conditions and qPCR procedure as described previously (16) to quantify the effect of Δ*ccrZ* on DNA replication initiation by measuring the origin to terminus ratio (*ori/ter*) **(Figure 1 and Table 1)**. In WT cells grown in rich medium (LB) at 37°C, we show that the *ori/ter* ratio is 3.59±0.28, a result consistent with multiple published reports (16, 34, 36). Cells carrying Δ*ccrZ* under-initiate showing an *ori/ter* ratio of 2.77±0.22 which is significantly lower than WT (p<0.05), supporting previous findings (16, 34). For comparison, we also quantified the *oriC/ter* ratio in cells lacking the DNA replication initiation negative regulator *yabA* and found a ratio of 5.28+0.17, a measurement significantly different from WT (p<0.001). When Δ*ccrZ* was combined with the *yabA* null allele, the ratio was 3.95±0.26, supporting prior work that under and over-initiation effects can balance to yield a near WT *oriC/ter* ratio (16, 34). As a control, we ectopically expressed *ccrZ* from the *ccrZ* promoter at *amyE* in a Δ*ccrZ* background [Δ*ccrZ*, *amyE*::*P_ccrZ_-ccrZ*] complementing the under-initiation effect with an *oriC/ter* ratio measurement of 4.26±0.09 **(Figure 1 and Table 1)**. We also tested the effect of Δ*ccrZ* and the *yabA* null on DNA replication initiation frequency with cells grown in minimal medium and showed the same overall trend as our observations described above for cells grown in rich medium **(Figure S1 and Table 1)**.

**Figure 1.**
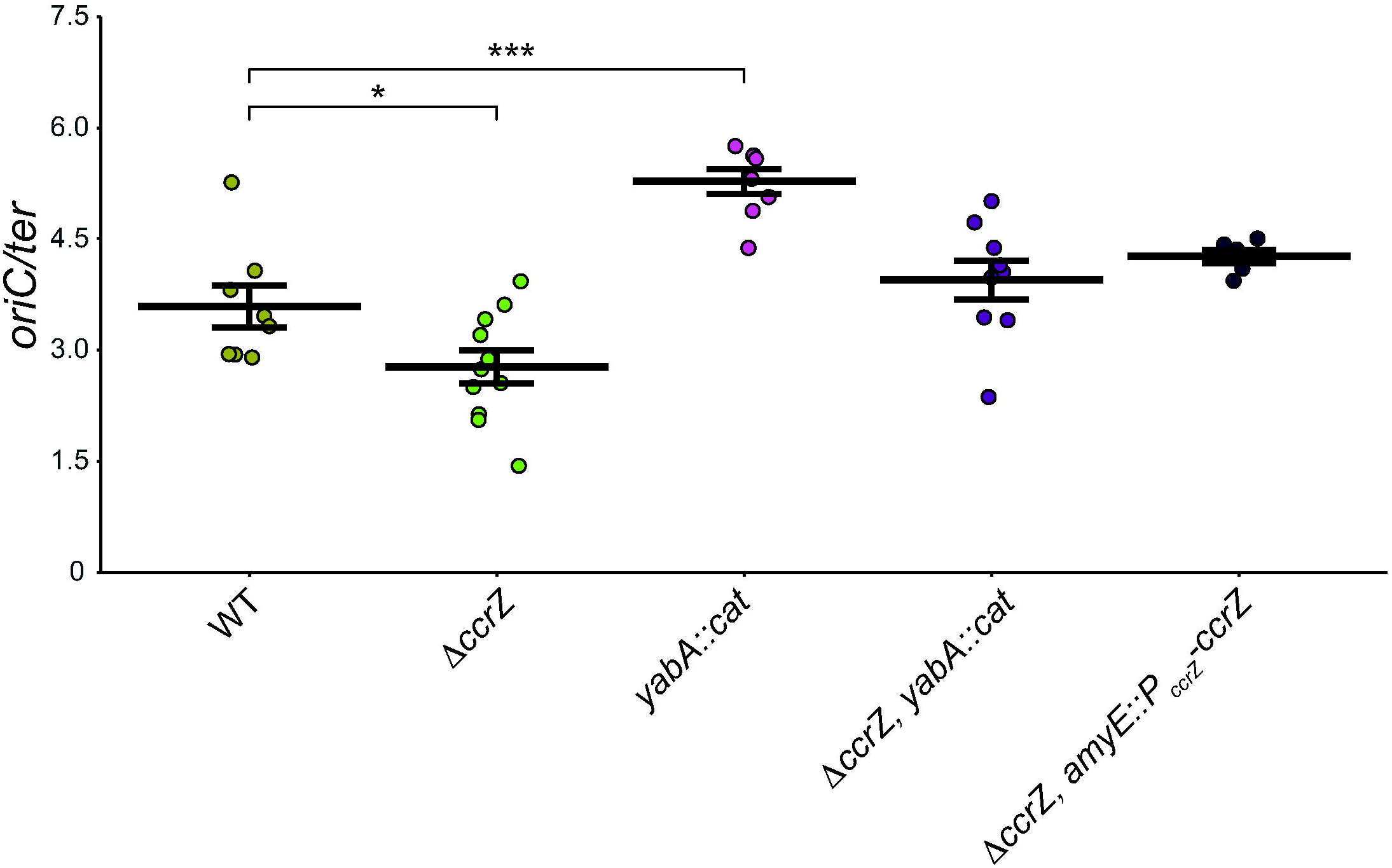
Establishing *oriC/ter* ratio of CcrZ expression under the *amyE* locus. Shown are qPCR-generated *oriC/ter* ratios for the *B. subtilis* strains indicated grown in LB medium at 37°C as described in Materials and Methods. Data points plotted with means represented by a horizontal bar and error bars representing standard error of the mean for at least six independent replicates. Statistical significance of increased or decreased ratios compared to WT assessed by two-tailed t-test, *p < 0.05, **p < 0.01, ***p < 0.001, ****p < 0.0001. The data shown here are also summarized in Table 1.

**Table 1.** Summary of *oriC/ter* ratios.

| **Genotype** | **Growth Medium** | ***oriC/ter*** |
| --- | --- | --- |
| WT | LB | 3.59±0.28 |
| $\Delta ccrZ$ | LB | 2.77±0.22 |
| *yabA::cat* | LB | 5.28±0.17 |
| $\Delta ccrZ$, *yabA::cat* | LB | 3.95±0.26 |
| $\Delta ccrZ$, *amyE::P<sub>ccrZ</sub>-ccrZ* | LB | 4.26±0.09 |
| $\Delta ccrZ$, *amyE::P<sub>xyl</sub>-ccrZ* | LB | 9.05±0.65 |
| $\Delta ccrZ$, *amyE::P<sub>xyl</sub>-ccrZ-D166A* | LB | 3.74±0.28 |
| $\Delta ccrZ$, *amyE::P<sub>ccrZ</sub>-ccrZ-D166A* | LB | 3.32±0.12 |
| *ccrZ<sup>+</sup>*, *amyE::P<sub>ccrZ</sub>-ccrZ-D166A* | LB | 2.74±0.22 |
| $\Delta ccrZ$, *amyE::P<sub>ccrZ</sub>-ccrZ-F47A* | LB | 2.76±0.22 |
| $\Delta ccrZ$, *amyE::P<sub>ccrZ</sub>-ccrZ-S103A* | LB | 4.39±0.36 |
| $\Delta ccrZ$, *amyE::P<sub>ccrZ</sub>-ccrZ-N171A* | LB | 3.31±0.05 |
| $\Delta ccrZ$, *amyE::P<sub>ccrZ</sub>-ccrZ-D184A* | LB | 3.26±0.08 |
| WT | S7<sub>50</sub>+ arabinose | 1.47±0.03 |
| $\Delta ccrZ$ | S7<sub>50</sub>+ arabinose | 1.19±0.06 |
| *yabA::cat* | S7<sub>50</sub>+ arabinose | 1.85±0.07 |
| $\Delta ccrZ$, *yabA::cat* | S7<sub>50</sub>+ arabinose | 1.63±0.10 |

To determine *ori/ter* ratios of over-initiating strains, we induced expression of *ccrZ* from a xylose inducible promoter from an ectopic chromosomal locus [Δ*ccrZ*, *amyE*::*P_xyl_-ccrZ*]. When *ccrZ* expression was induced, we quantified an *ori/ter* ratio of 9.05±0.65, 2.5-fold higher than WT and more than 3-fold greater than Δ*ccrZ* **(Figure 2 and Table 1)**. Our qPCR results are similar to genome re-sequencing experiments showing that overproduction of CcrZ results in an elevated *oriC/ter* ratio (35). Collectively, our experiments show that overproduction of CcrZ results in a striking increase in replication initiation, a more severe initiation phenotype than our measurement of 5.28+0.17 for cells lacking *yabA*. We conclude that overproduction of *ccrZ* results in a substantial increase in replication initiation activity.

**Figure 2.**
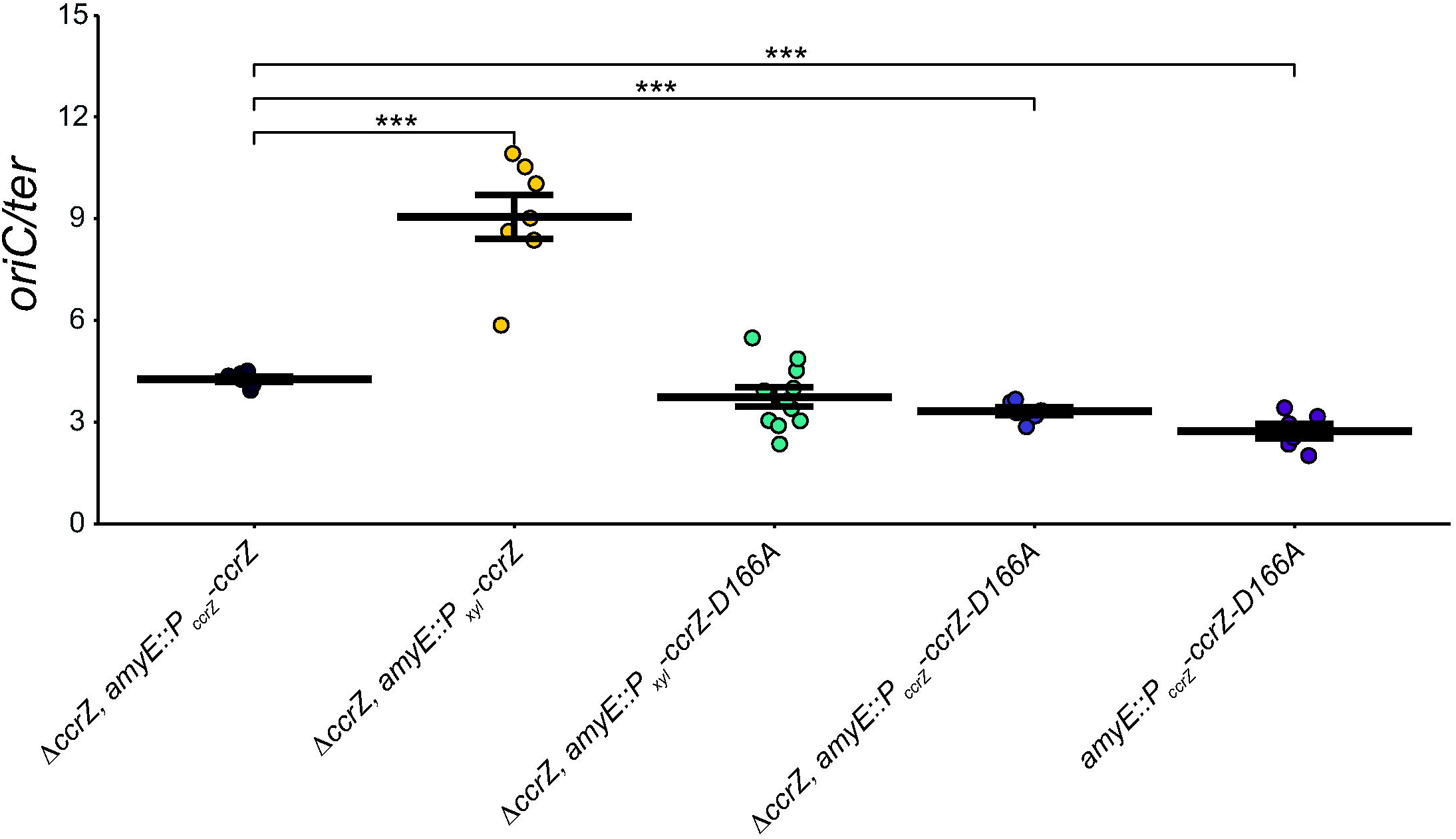
*ccrZ* and *ccrZ* mutant D166A under xylose-inducible promoters lead to increased *oriC/ter* ratios. Strains were grown in LB medium and 0.25% xylose in LB medium used for all strains with a xylose-inducible promoter. The *oriC/ter* ratios generated by qPCR are shown as individual data points. The crossbar represents the mean and error bars shown indicate standard error of the mean for at least six independent replicates. Statistical significance assessed by two-tailed t-test compared to Δ *crZ, amyE::P_ccrZ_-ccrZ*, *p < 0.05, **p < 0.01, ***p < 0.001, ****p < 0.0001. The data shown here are also summarized in Table 1.

### Mutations in *ccrZ* under-initiate DNA replication

Based on the crystal structure of CcrZ (35), we tested several mutants for replication initiation activity *in vivo*. CcrZ(D166A) has been purified and shown to be defective in kinase activity (35). We overproduced *ccrZ*(*D166A*) in a Δ*ccrZ* background and measured an *oriC/ter* ratio of 3.74±0.28, a measurement that is the same as WT but higher than Δ*ccrZ* **(Figure 2 and Table 1)**. Since CcrZ has been shown to interact with DnaA and DnaB, we hypothesize that the slight initiation stimulation is caused by interactions between kinase-inactive CcrZ and DnaA or DnaB. To test this, we complemented Δ*ccrZ* with *ccrZ*(*D166A*) expressed from its own promoter from the *amyE* ectopic locus and measured a lower *oriC/ter* ratio of 3.32±0.12. When *ccrZ*(*D166A*) was expressed ectopically with the WT *ccrZ* allele intact we measure a ratio of 2.76+0.22 indicating that *ccrZ(D166A)* is dominant negative to WT *ccrZ* **(Figure 1, 2 and Table 1)**. With these results we suggest that CcrZ(D166A) is indeed defective in replication initiation. However, when this variant is overproduced, it is able to partially compensate for the loss of kinase activity through stimulatory interactions with DnaA and/or DnaB resulting in an *oriC/ter* ratio closer to WT levels **(see Discussion)**.

Since we found that overproduction of CcrZ(D166A) yields a WT ratio of *oriC/ter*, we tested *ccrZ F47A, S103A, N171A* and *D184A* variants each driven by the *ccrZ* promoter in cells with the *ccrZ* deletion allele **(Figure 3 and Table 1)**. Residues N171 and D184 are hypothesized to coordinate Mg^2+^ based on the crystal structure (35). F47 provides a hydrophobic group at the interlobal cleft, hypothesized to maintain proper distance between the N and C terminal lobes of the enzyme (35). S103 is predicted to be important for forming an interface between the last two alpha helices on the C-terminus (35). The *ccrZ* mutants *N171A* and *D184A* supported replication activity greater than Δ*ccrZ* but less than WT, indicating these mutants are indeed compromised for CcrZ activity *in vivo.* The difference between WT and N171A and D184A were significant with p<0.0001 **(Figure 3)**. We found that F47A was defective in supporting replication initiation with an *oriC/ter* ratio of 2.76±0.22, yielding a ratio indistinguishable from Δ*ccrZ*. We found that S103A was active in replication initiation yielding a ratio of 4.39±0.36. With our results we suggest that F47 provides a critical role for enzyme activity in maintaining the proper distance between the N and C-terminal lobes of the enzyme, while S103A results in a WT replication initiation frequency. All defective protein variants tested have been shown to accumulate to WT levels *in vivo* (35), indicating that the defects we observe in replication initiation are not due to low levels of expression or accumulation of CcrZ variants in the cell.

**Figure 3.**
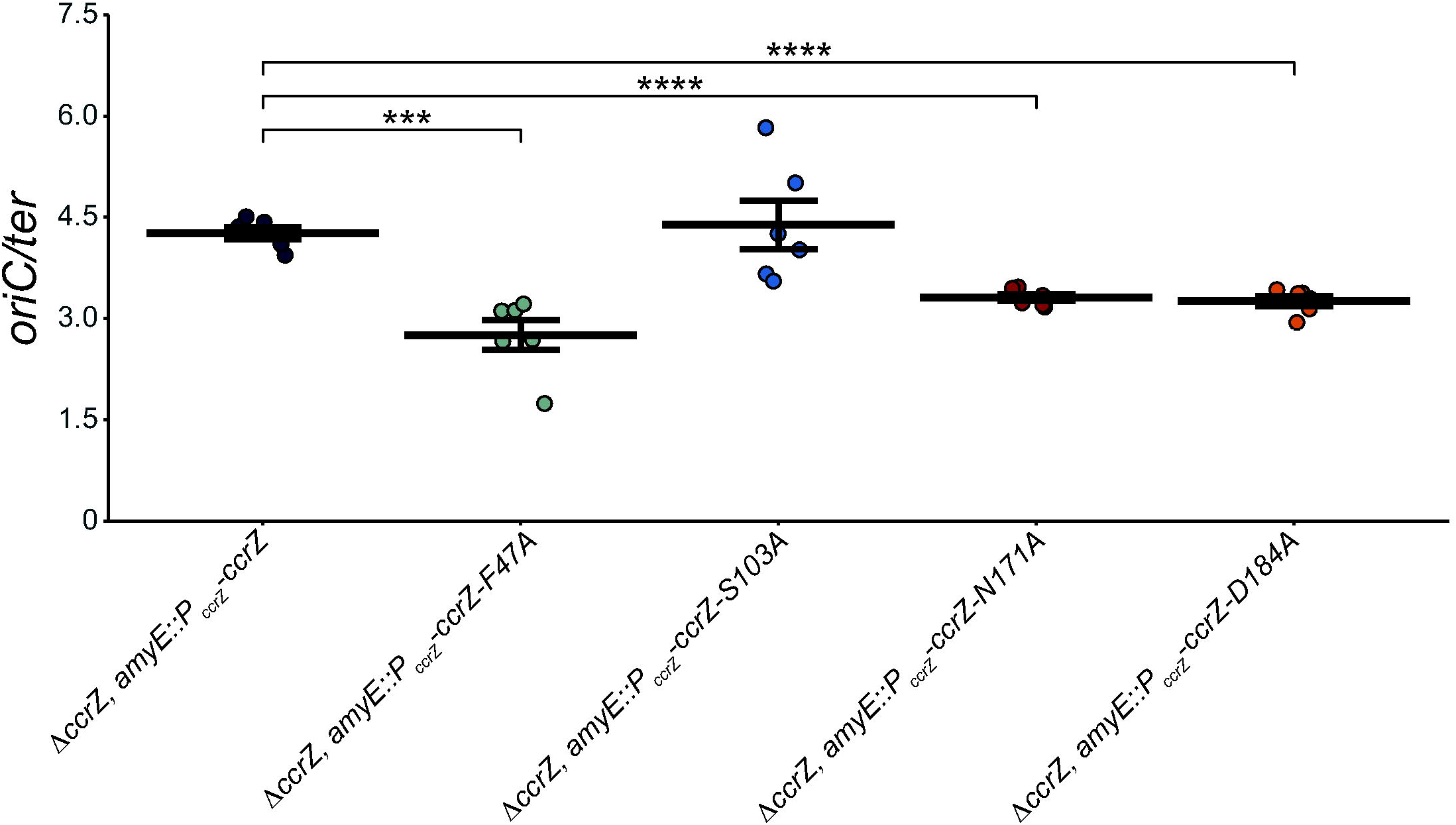
Mutations that disrupt CcrZ function have lowered *oriC/ter* ratios. Various strains carrying *ccrZ* mutants were grown in LB media with *oriC/ter* ratios assessed by qPCR. The individual data points are shown. The horizontal bars indicate the mean while the error bars represent standard error of the mean for each strain. The data shown are from six independent replicates. Statistical significance compared to Δ*ccrZ, amyE::P_ccrZ_-ccrZ* assessed by two-tailed t-test, *p < 0.05, **p < 0.01, ***p < 0.001, ****p < 0.0001. The data shown here are also summarized in Table 1.

In our prior work, we showed that cells with a *yabA* null allele or certain *ccrZ* alleles conferred sensitivity to DNA damage (35). We show above that *ccrZ F47A*, *D166A, N171A*, and *D184A* yield similar DNA replication activity as Δ*ccrZ* while *S103A* showed replication initiation activity similar to WT. We therefore tested each *ccrZ* allele for sensitivity to mitomycin C (MMC). We show that *F47A, D166A, N171A*, and *D184A* each confer sensitivity to MMC to a similar extent as Δ*ccrZ,* while *S103A* and the complementing strain show WT growth in the presence of MMC, confirming previous observations **(Figure 4)** (35). Collectively, these experiments show that under-initiation of DNA replication is correlated with increased sensitivity to genotoxic stress. We speculate that because Δ*ccrZ* cells under-replicate, there is less homologous DNA available for recombination to repair MMC-induced damage, resulting in greater growth interference in the presence of DNA damage.

**Figure 4.**
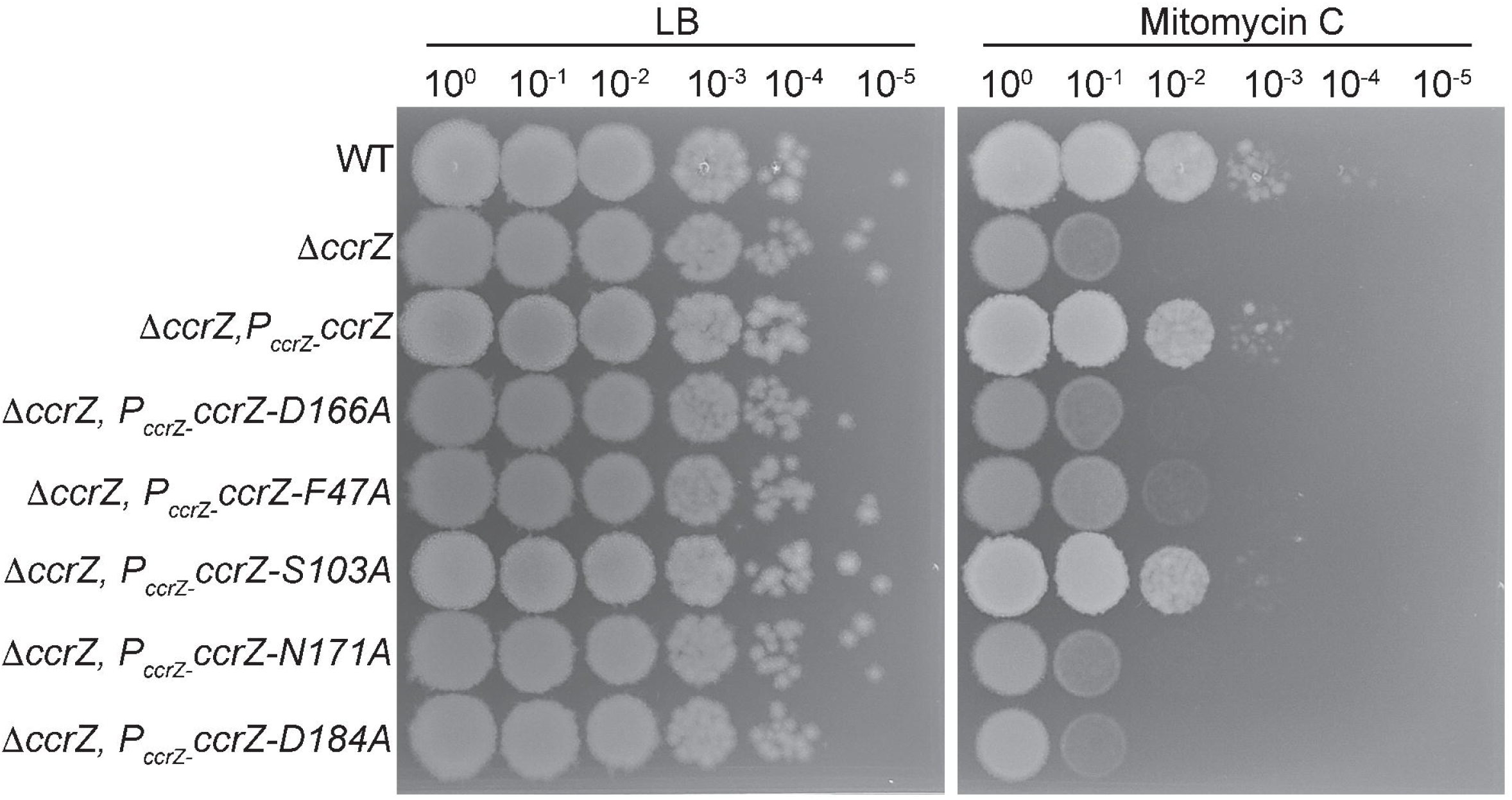
*ccrZ* alleles defective in replication initiation confer sensitivity to DNA damage. Shown is a spot titer assay with the indicated strains spotted onto LB or LB with 25ng/ml mitomycin C (MMC). Each *ccrZ* allele is expressed from the *amyE* locus undercontrol of the native *ccrZ* promoter as described (35).

### Over-initiation of DNA replication leads to replication fork stress

Prior work in *E. coli* showed that over-initiation of DNA replication through overproduction of DnaA or using the DnaAcos hyperactive variant resulted in replication fork stalling as determined by tracking replication fork progress using microarrays (19). When CcrZ was overproduced in *B. subtilis,* whole genome resequencing showed an increase in origin DNA, although the slope and marker frequency from the origin to the terminus provided no indication of replication fork stalling (35). Even though the slope of fork movement *in vivo* following CcrZ overexpression was inconsistent with fork stalling, we reasoned that cells could still experience replication fork stress without forks accumulating near the origin. Replication fork stress occurs when the fork encounters lesions, protein impediments or other problems that generate excess ssDNA, leading to RecA binding (37, 38). Excess ssDNA bound by RecA, leads to the formation of RecA-GFP foci at the replication fork in *B. subtilis* cells (37–39). Therefore, we used RecA-GFP as a proxy for replication fork stress in single *B. subtilis* cells following over-initiation of DNA replication by disrupting *yabA* or by overproducing *ccrZ* **(Figure 5 and Table 2)**. In WT cells, RecA-GFP forms foci in ∼12% (n=2550) of cells, supporting prior findings (37–39). Cells disrupted for *yabA* or overproducing *ccrZ* showed RecA-GFP foci in ∼23.5 (n=1941) and ∼37.4% (n=891) of cells, respectively. In contrast, cultures of cells with Δ*ccrZ* only showed RecA-GFP foci in ∼8.4% (n=2674) of cells, significantly lower than WT cells. These results show that elevated initiation of DNA replication leads to excess ssDNA that causes RecA-GFP to organize into foci, while under-initiation of DNA replication in Δ*ccrZ* has less DNA and less RecA-GFP foci suggesting that homologous recombination is reduced **(see Discussion)**.

**Figure 5.**
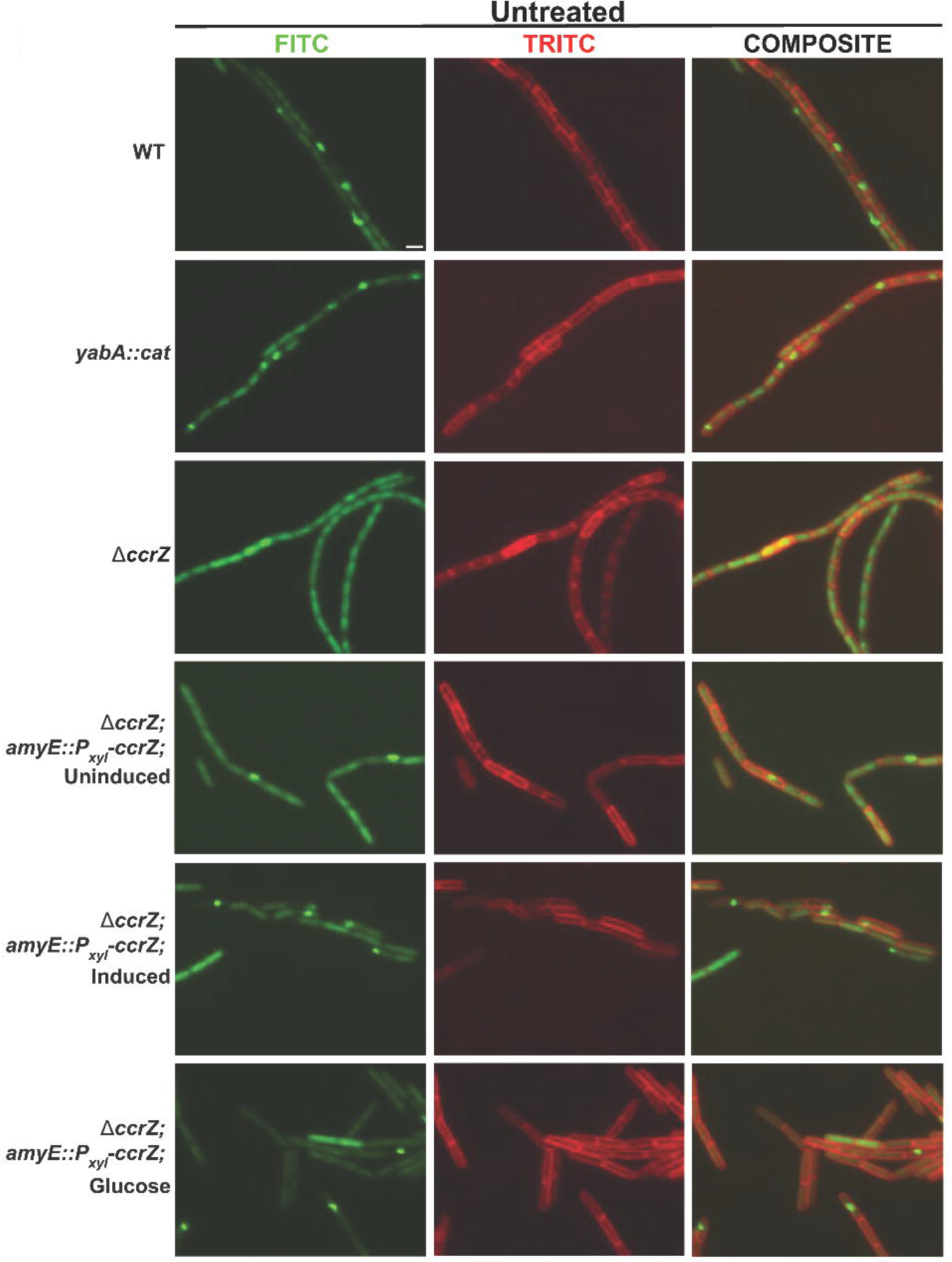
RecA-GFP foci are elevated in cells with increased DNA replication initiation. Shown are representative images of cells with the indicated genotypes. The left most column is RecA-GFP, followed by the middle column membrane stained with FM 4-64 and the merge of the GFP and FM 4-64 channels (composite) to the right.

**Table 2.** Percent of cells with RecA-GFP foci.

| Strain Name | Growth Medium | Percent of cells<br>with foci<br>(Untreated) | Percent of cells<br>with foci<br>(Mitomycin C) |
| --- | --- | --- | --- |
| WT *recA-gfp* | S7<sub>50</sub> glucose | 12.3 ± 1.0<br>(n = 2550) | 43.5 ± 0.6<br>(n = 1365) |
| *yabA::cat; recA-gfp* | S7<sub>50</sub> glucose | 23.5 ± 1.2<br>(n = 1941) | 78.5 ± 2.1<br>(n = 1765) |
| *ΔccrZ; amyE::P<sub>xyl</sub>-*<br>*ccrZ; recA-gfp* | S7<sub>50</sub> arabinose<br>(uninduced) | 3.9±0.5<br>(n=1333) | 38.0±7.0<br>(n=987) |
| *ΔccrZ; amyE::P<sub>xyl</sub>-*<br>*ccrZ; recA-gfp* | S7<sub>50</sub> arabinose<br>(induced) | 37.4 ± 5.4<br>(n = 891) | 63.1 ± 4.9<br>(n = 773) |
| *ΔccrZ; amyE::P<sub>xyl</sub>-*<br>*ccrZ; recA-gfp* | S7<sub>50</sub> glucose | 9.3 ± 1.6<br>(n = 1045) | 17.9 ± 2.3<br>(n = 998) |
| *ΔccrZ; recA-gfp* | S7<sub>50</sub> glucose | 8.4 ± 0.4<br>(n = 2674) | 33.0 ± 0.5<br>(n = 1135) |

Relationships between the untreated and treated condition of each strain were found to be statistically significant by a Welch two-sample t-test. Standard error was calculated using standard deviation and sample size. S7<sub>50</sub> is the minimal medium used for growth of each culture. *ccrZ* expression is induced with the addition of 1% xylose to S7<sub>50</sub> with arabinose as a carbon source.

We asked if cells challenged with DNA damage showed elevated RecA-GFP foci following over-initiation of DNA replication. We reasoned that RecA-GFP foci formation would become exacerbated in damaged cells that also over-initiate DNA replication. We challenged WT cells with a concentration of MMC that would lead to elevated RecA-GFP foci without saturating the response **(Figure S2)**. For WT cells, we scored RecA-GFP foci in ∼43.5% (n=1365) of cells following the addition of MMC. Cells deficient for *yabA* or overproducing *ccrZ* showed foci in ∼78.5% (n=1765) and ∼63.1% (n=773) of cells respectively, while cells with Δ*ccrZ* had RecA-GFP foci formation in ∼33% of cells (n=1135), lower than that of WT **(Figure S3 and Table 2)**. We wish to note that otherwise untreated cells overproducing CcrZ showed foci in ∼37% of cells while WT cells challenged with MMC formed foci in ∼43% percent of cells. These results show that unregulated replication initiation results in a level of genome instability very similar to cells treated with an exogenous source of DNA damage. We conclude that over-initiation of DNA replication leads to changes in the formation of replication fork problems which are further exacerbated by the presence of damaged template DNA. Further, RecA-GFP foci are closely correlated with DNA replication initiation frequency. Cells that under initiate replication have fewer foci than WT cells and cells that over-initiate have more foci compared with WT. Our results are similar to results found in *E. coli,* showing that the percent of cells with RecA-GFP foci correlates with growth rate (40).

Since it has been shown in eukaryotes that over-initiation of replication leads to detrimental conflicts between DNA replication and transcription (27), we asked if the correlation between over-initiation and RecA-GFP foci formation was caused by replication forks encountering R-loops. In *B. subtilis* RNase HIII is responsible for cleaving R-loops *in vitro* and *in vivo* (41, 42). Therefore, we overexpressed RNase HIII (*rnhC*) in cells with the *yabA* null allele and plated on MMC. If replication forks in over-initiating cells encountered R-loops leading to genome instability, we would expect overexpression of RNase HIII to rescue or at least partially rescue the sensitivity of *yabA* null cells to DNA damage. We observed no rescue of *yabA* null cells when RNase HIII was overexpressed **(Figure S4)**. We did, however, find better survival of otherwise WT cells when *rnhC* was overproduced, **(Figure S4)** suggesting that R-loops are detrimental in WT cells that are challenged with DNA damage. We do however show that *yabA* null cells overproducing *ccrZ* caused an increased sensitivity to MMC, further demonstrating the connection between replication initiation frequency and sensitivity to DNA damage **(Figure S5)**.

## DISCUSSION

Here we investigated the effect of under-initiation and over-initiation on genome integrity using *B. subtilis*. We examined *ccrZ* mutants that cause under-initiation and overproduced CcrZ and used a *yabA* null allele to cause over-initiation. We find that cells under-initiating DNA replication have few RecA-GFP foci indicating that these cells incur less replication fork stress. We also find that cells under-initiating DNA replication are sensitive to DNA damage caused by MMC. We suggest that under-initiating cells have less homologous DNA that can be used during homologous recombination leading to the MMC sensitive growth phenotype. Our finding also correlates with fewer RecA-GFP foci in cells that under initiate suggesting that either forks experience less problems that require RecA and/or recombination in general is reduced in cells that under-initiate replication.

In cells over-initiating DNA replication, we find a substantial increase in both the sensitivity to DNA damage and an increase in RecA-GFP localization. Further, another important finding from our work is that over-initiation of replication leads to nearly the same percentage of cells with RecA-GFP foci as WT cells directly damaged with the crosslinking agent MMC. This result suggests that over-initiation has a strong negative effect on genome integrity. We conclude that an important feature of regulating DNA replication initiation frequency is to maintain genome stability as under or over-initiation results in sensitivity to genotoxic stress and increased fork stress in cells that over initiate.

CcrZ is a recently identified positive regulator of DNA replication initiation in *B. subtilis* (34, 35). CcrZ is a kinase and it has been shown to interact with DnaA and DnaB through a bacterial two hybrid assay (35). We measure *oriC/ter* ratios using *ccrZ* alleles under control of the native promoter or following induced expression. We show that CcrZ(D166A), which is known to be defective for kinase activity yields a WT *oriC/ter* ratios when overexpressed, but not when *ccrZ(D166A)* is expressed from its own promoter. We suggest that the WT ratios measured in cells overproducing CcrZ is due to weak stimulation of initiation through interaction with the *B. subtilis* replication initiation proteins. The replication initiation stimulation observed with CcrZ(D166A) is far lower than when WT CcrZ is overproduced. With this data, we conclude that kinase activity from CcrZ provides the most important contribution to stimulating replication initiation activity with interaction between CcrZ, DnaA and DnaB providing some positive effect.

It has been clearly established in eukaryotes that under and over-initiation of DNA replication leads to R-loop mediated replication fork stress and DNA damage at fragile sites [for review (25)]. We overexpressed *B. subtilis* RNase HIII to test if R-loop-induced replication stress was occurring in cells undergoing over-initiation. We did not observe any rescue suggesting that R-loops either do not pose a major threat to genome stability in over-initiating cells or RNase HIII is inefficient at clearing R-loops encountered by replication forks under the conditions tested here. Since ectopic expression of RNase HIII yielded more survival to WT cells plated on DNA damage, we speculate that R-loops are likely to threaten genome integrity during over-initiation. However, we further speculate that cells over-initiating DNA replication and contending with DNA damage are compromised to the point where R-loop removal by RNase HIII is unable to restore even partial growth.

Overall, we conclude that the proper regulation of DNA replication initiation is central to limiting replication fork stress and maintaining genome stability in bacterial cells that undergo multi-fork replication.

## Materials and Methods

### Media and Growth Conditions

All strains and primers used in this study are described in supplemental **Tables S1** and **S2**. All strains are derivatives of PY79 (43) and were constructed using standard procedures (44). Cultures for *oriC/ter* ratio generation were grown at 37°C in a water bath with shaking at 250 rpm. Lysogeny broth (LB medium: 10 g/L NaCl, 10 g/L tryptone, and 5 g/L yeast extract) or S7_50_ defined minimal medium with 2% glucose as described previously (35). In LB or S7_50,_ xylose-inducible promoters were induced with 0.25% xylose. Antibiotics were used at the following final concentrations: spectinomycin (100 μ mL), kanamycin (50 μ mL) and chloramphenicol (5 µg/mL).

### Microscopy

Strains were grown on LB agar plates with the appropriate antibiotics and incubated at 37°C overnight. Cells were grown in S7_50_ minimal medium with 2% glucose or, 1% arabinose as a carbon source. 12 mL S7_50_ medium was added to a 125 mL Erlenmeyer flask and placed in a 30°C shaking water bath incubator for approximately 5-7 minutes. Then, 1.5 mL of the pre-warmed media was washed over the agar plate and the optical density of the starting culture was measured between 0.080 and 0.1). The flask was wrapped tightly in aluminum foil, returned to the incubator, and grown at 30°C at 250 rpm until an OD_600_ value of 0.5-0.7 was reached. In order to assess the impact of DNA damage on the prevalence of RecA-GFP repair centers, cells were treated with the DNA crosslinking agent mitomycin C (MMC). 5 mL of the culture was transferred to a 50 mL Erlenmeyer flask where 2.5 ng/mL of MMC was added. The flask was returned to the incubator and grown for 30 minutes prior to the start of imaging.

After cultures reached an OD_600_ of 0.5-0.7 or following treatment with MMC, 300 μ L (2 µg/mL) of FM4-64 membrane dye and gently agitated to incorporate the dye. 1% agarose was melted into 1XS7_50_ before being pipetted onto a standard 15-well glass slide. Cells were pipetted onto the agarose pads and allowed to settle for 7-10 minutes. Excess liquid was then aspirated from each well. A coverslip was set on top of the slide and adhered for approximately 2 minutes. Cells were imaged using an Olympus BX61 microscope under TRITC (exposure set between 200-500ms) and FITC (exposure set between 400-800ms) filters as described (37, 39, 45, 46). Shapiro-Wilk tests were performed to assess if microscopy data were normally distributed. The tests suggested that the data collected met the requirements for a normal distribution, thus, Welch’s t-tests were used to assess statistical significance between strains. In order to analyze data within a single strain and across conditions, dependent t-tests were used.

### oriC/ter ratio generation via qPCR

In defined minimal medium or LB medium, cultures were started at an initial OD_600_ = 0.05 by diluting a plate wash into the respective medium. When cultures reached an OD_600_ between 0.2-0.4, they were back diluted to OD_600_ = 0.05, grown to OD_600_ of 0.2-0.4 again, and immediately arrested in ice-cold methanol in a 1:1 ratio of culture to methanol. The resulting mixture of methanol and culture were pelleted at 7197 ×□ for 5 minutes. Genomic DNA was purified as described below. qPCR was used to generate origin (*oriC*) to terminus (*ter*) ratios, quantifying the relative amounts of each via the Pfaffl method as described (16, 47). qPCR was performed with 20 μ reactions containing 10 μ SsoAdvanced Universal SYBR Green Supermix, 250 nM of each primer, and 4 ng of gDNA. StepOnePlus™ Real-Time PCR System used for all qPCR runs. Primers targeting origin region were oTTR1 (5′ TTGCCGCAGATTGAAGAG-3 ) and oTTR2 (5′ AGGTGGACACTGCAAATAC-3 ) while primers targeting terminus region -′were oTTR3 (5’-CGCGCTGACTCTGATATTATG-3’) and oTTR4 (5′-CAAAGAGGAGCTGCTGTAAC-3′), primers are also listed in Table S2 and were used in the following manuscript (16). *oriC/ter* ratios were generated in reference to a temperature sensitive mutant, *dnaB134* (TTR1), grown identically to other samples followed by incubation at its non-permissive temperature of 45°C for two additional hours in a still bath, allowing for replication fork runout and resulting in a 1:1 ratio of origin to terminus DNA.

### Genomic DNA purification

Cell pellets from 25 mL cultures were resuspended in 200 μ of lysis buffer (50 mM Tris-HCl [pH 8], 10L M EDTA [pH 8], 1% Triton X-100, 0.5 mg/ml RNase A, 5 mg/ml lysozyme) followed by incubation for 30 min at 37°C. Next, 40 μ of 10% SDS and Proteinase K were added and mixed by pipette. Samples were subsequently incubated at 55°C for 55 minutes. 600 μ PB buffer (5 M guanidine-HCl, 30% [vol/vol] isopropanol) was added, gently mixed, and the sample was applied to EconoSpin^TM^ spin column for DNA and centrifuged for 1 min at 12,000□×□*g*. 500 μl PB was used to wash the column and centrifuged for the same speed and time as prior. 500 μl of PE (10 mM Tris-HCl [pH 7.5], 80% [vol/vol] ethanol) was added for a final wash, centrifuged at 12,000 ×□*g* for 1 minute, followed by an additional centrifugation of the dry column for a minute at 12,000□×□*g*. 10 mM Tris-HCl [pH 8] was used to elute the DNA, centrifuging at 14,000 *x g* for 1 minute. The eluant was reloaded onto the column and centrifuged at 14,000 *x g* for 1 minute.

## Supporting information

All supplemental figures and tables.

## SUPPLEMENTAL MATERIALS

**Figure S1. Relative *oriC/ter* ratios of cultures grown in S7_50_ minimal medium with 1% arabinose.**

Shown are qPCR-generated *oriC/ter* ratios for *B. subtilis* strains grown in S7_50_ minimal medium with 1% arabinose at 37°C. Data points are plotted along with their means repressented by the horizontal bar and error bars represent the standard error of the mean for at least six biological replicates. Statistically significant increase or decrease in ratios assessed by two-tailed t-test in relation to WT, *p < 0.05, **p < 0.01, ***p < 0.001, ****p < 0.0001.

**Figure S2. Organization of RecA-GFP into foci in response to mitomycin C concentration.** The proportion of cells with RecA-GFP foci were determined in response to the concentration of mitomycin C (MMC) added to the culture medium. Cells were challenged with the indicated concentration of MMC for 30 minutes prior to imaging. The percent of cells with foci and the number of cells scored (n) are as follows: untreated (12.5%, n=594); 2.5 ng/mL (43.3%, n=531); 5 ng/mL (69.9%, n=596) 10 ng/mL (73.2%, n=504); 25 ng/mL (84.1%, n=584). Based on this data 2.5 ng/mL was chosen as the appropriate concentration of MMC to test for RecA-GFP foci in cells over and under-initiating DNA replication.

**Figure S3. Cells during over-initiation of DNA replication have more RecA-GFP foci.**

Shown are representative micrographs of cells with the indicated genotypes grown normally in S7_50_ minimal medium and challenged with 2.5 ng/ml mitomycin C for 30 minutes prior to imaging. The left most column is RecA-GFP, followed by the membrane stained with FM 4-64 followed by the merge of the GFP and FM 4-64 channels (composite).

**Figure S4. Overexpression of RNase HIII does not alleviate growth interference caused by over-initiation of DNA replication.** Shown are spot titer plates of the indicated strains on LB or LB with 25 ng/ml mitomycin C. Expression of *rnhC* (RNase HIII) is driven by the IPTG regulated P_spac_ promoter

**Figure S5. Expression of *ccrZ* in *yabA* defective cells results in growth interference and sensitivity to DNA damage.** Shown are spot titers on the indicated medium with increasing 10-fold serial dilutions. Expression of CcrZ is driven by xylose using the P_xyl_ promoter with MMC added to 25 ng/ml.

**Table S1. Strains used in this study.**

**Table S2. Primers used in this study.**

## DATA AVAILABILITY

The raw data for the qPCR experiments underlying the data in Table 1, Figures 1-3 and Figure S1 are deposited at https://figshare.com/browse https://doi.org/10.6084/m9.figshare.25938388.v1

## ACKNOWLEDGEMENTS

We thank members of the Simmons lab for their helpful contributions during the development of this work. This work was funded by National Institutes of Health grant R35GM131772 to LAS. KJW was supported by a predoctoral fellowship from the National Science Foundation (award DEG 1256260) and by the NIH Cellular Biotechnology Training Program (award T32 GM008353).

