## Supplementary material for "DNA replication initiation timing is important for maintaining genome integrity": All supplemental figures and tables.

Short title: DNA replication initiation impacts genome integrity

**Keywords:** CcrZ, *Bacillus subtilis,* DnaA, DNA replication, RecA

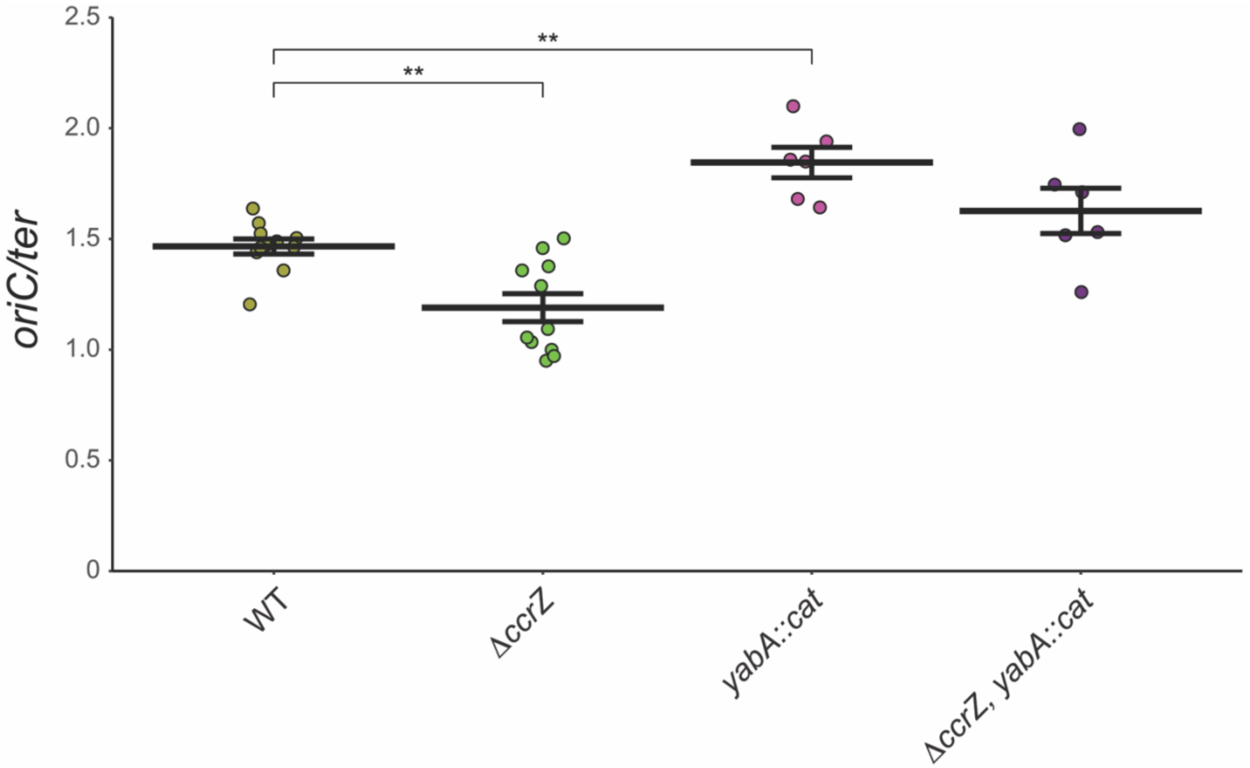


**Figure S1. Relative *oriC/ter* ratios of cultures grown in $S7_{50}$ minimal medium with 1% arabinose.**

Shown are qPCR-generated *oriC/ter* ratios for *B. subtilis* strains grown in $S7_{50}$ minimal medium with 1% arabinose at 37°C. Data points are plotted along with their means represented by the horizontal bar and error bars represent the standard error of the mean for at least six biological replicates. Statistically significant increase or decrease in ratios assessed by two-tailed t-test in relation to WT, *$p < 0.05$, **$p < 0.01$, ***$p < 0.001$, ****$p < 0.0001$.

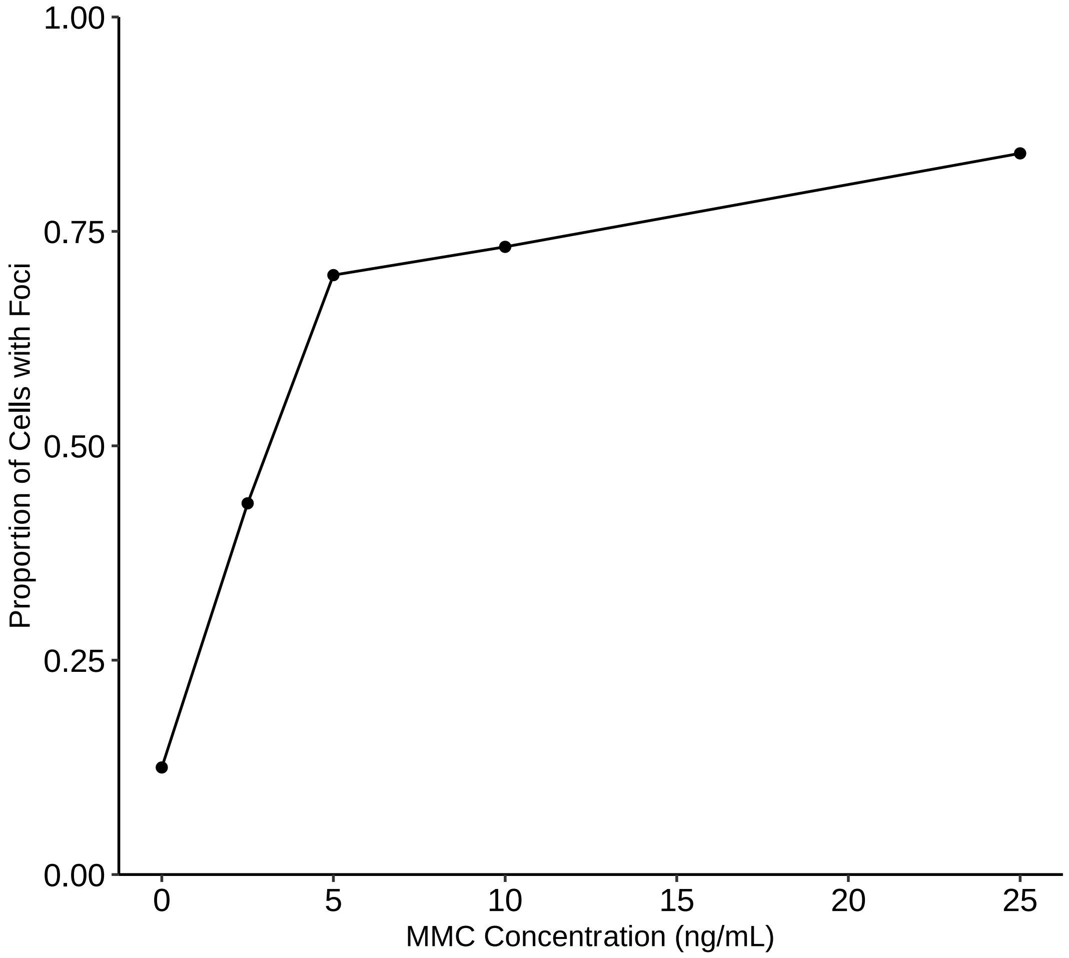


**Figure S2. Organization of RecA-GFP into foci in response to mitomycin C concentration.** The proportion of cells with RecA-GFP foci were determined in response to the concentration of mitomycin C (MMC) added to the culture medium. Cells were challenged with the indicated concentration of MMC for 30 minutes prior to imaging. The percent of cells with foci and the number of cells scored (n) are as follows: untreated (12.5%, n=594); 2.5 ng/mL (43.3%, n=531); 5 ng/mL (69.9%, n=596) 10 ng/mL (73.2%, n=504); 25 ng/mL (84.1%, n=584). Based on this data 2.5 ng/mL was chosen as the appropriate concentration of MMC to test for RecA-GFP foci in cells over and under-initiating DNA replication.

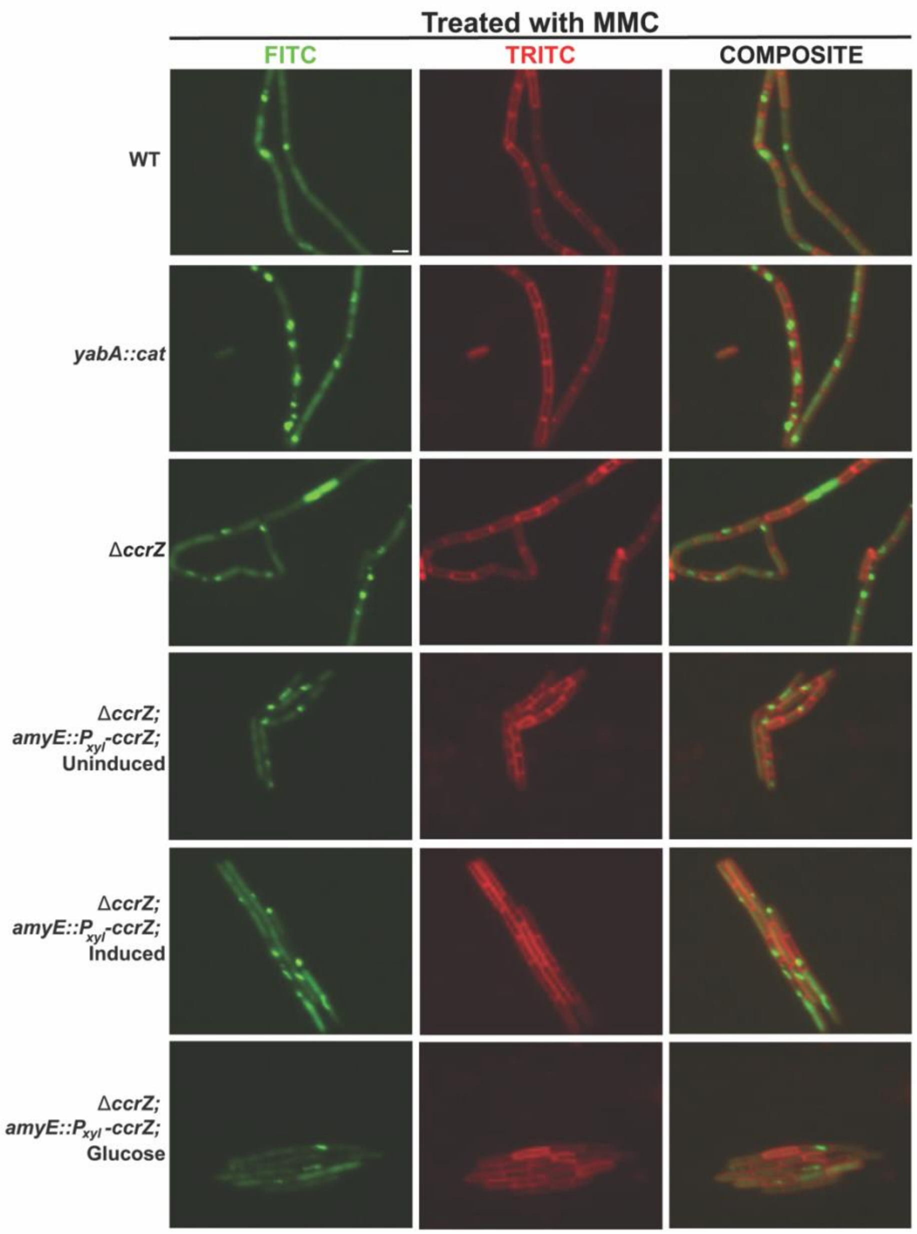


**Figure S3. Cells during over-initiation of DNA replication have more RecA-GFP foci.**

Shown are representative micrographs of cells with the indicated genotypes grown normally in $S7_{50}$ minimal medium and challenged with 2.5 ng/ml mitomycin C for 30 minutes prior to imaging. The left most column is RecA-GFP, followed by the membrane stained with FM 4-64 followed by the merge of the GFP and FM 4-64 channels (composite).

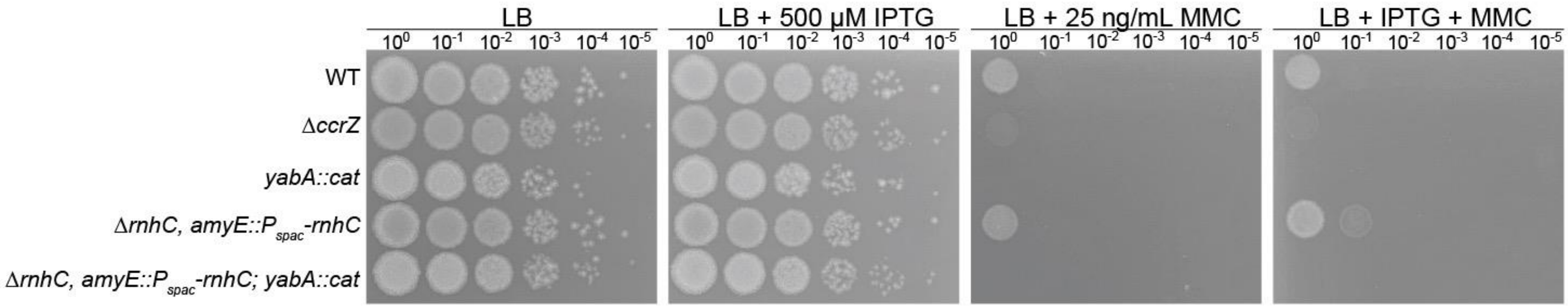


**Figure S4. Overexpression of RNase HIII does not alleviate growth interference caused by over-initiation of DNA replication.** Shown are spot titer plates of the indicated strains on LB or LB with 25 ng/ml mitomycin C. Expression of *rnhC* (RNase HIII) is driven by the IPTG regulated $P_{spac}$ promoter.

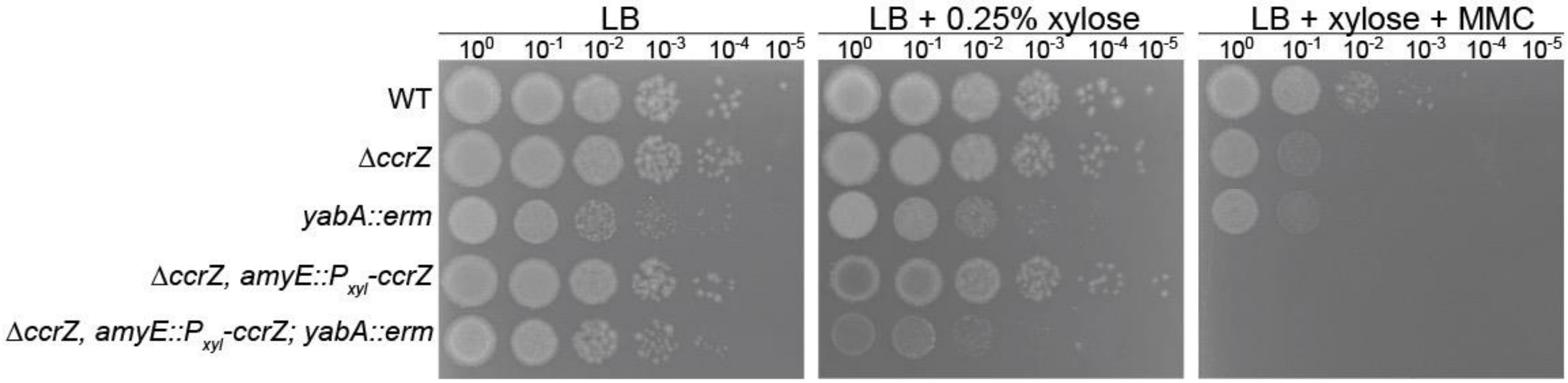


**Figure S5. Expression of *ccrZ* in *yabA* defective cells results in growth interference and sensitivity to DNA damage.** Shown are spot titers on the indicated medium with increasing 10-fold serial dilutions. Expression of CcrZ is driven by xylose using the $P_{xyl}$ promoter with MMC added to 25 ng/ml.

**Table S1. Strains used in this study.**

| Strain | Relevant Genotype | Source |
|---|---|---|
| PY79 | WT, prototroph SPβ° | (1) |
| TTR1 | *dnaB134(ts)*; *zhb83::Tn917* ($Cm^r$) | (2) |
| TTR4 | Δ*ccrZ* | (3) |
| TTR7 | Δ*ccrZ*, *amyE*::$P_{xyl}$-*ccrZ* ($Cm^r$) | (3) |
| TTR10 | ΔccrZ, *amyE*::$P_{xyl}$-*ccrZ*-D166A ($Cm^r$) | (3) |
| TTR19 | *yabA::cat* ($Cm^r$) | (4) |
| TTR22 | Δ*ccrZ, yabA::cat* ($Cm^r$) | (3) |
| TTR28 | Δ*ccrZ*, *amyE*::$P_{ccrZ}$-*ccrZ-D166A* ($Cm^r$) | (3) |
| TTR31 | Δ*ccrZ*, *amyE*::$P_{ccrZ}$-*ccrZ-F47A* ($Cm^r$) | (3) |
| TTR34 | Δ*ccrZ*, *amyE*::$P_{ccrZ}$-*ccrZ-S103A* ($Cm^r$) | (3) |
| TTR37 | Δ*ccrZ*, *amyE*::$P_{ccrZ}$-*ccrZ-N171A* ($Cm^r$) | (3) |
| TTR40 | Δ*ccrZ*, *amyE*::$P_{ccrZ}$-*ccrZ-D184A* ($Cm^r$) | (3) |
| TTR43 | Δ*ccrZ,amyE::P*$_{ccrZ}$-*ccrZ* (Cmr) | (3) |
| TTR46 | Δ*ccrZ, amyE::P*$_{ccrZ}$-*ccrZ-D166A* ($Cm^r$) | (3) |
| AHK1 | *recA::recA-gfp* ($Spc^r$) | (5) |
| AHK6 | *yabA::cat, recA::recA-gfp* ($Spc^r$) | This work |
| AHK18 | Δ*ccrZ, recA::recA-gfp* ($Spc^r$) | KJW868 |
| AHK16 | Δ*ccrZ, amyE*::$P_{xyl}$-*ccrZ* ($Cm^r$), *recA::recA-gfp* ($Spc^r$) | KJW870 |
| AHK20 | Δ*rnhC*, amyE::$P_{spac}$-*rnhC*, *yabA::cat* | This work |
| AHK26 | Δ*ccrZ, amyE*::$P_{xyl}$-*ccrZ* ($Cm^r$), *yabA::erm* | This work |

**Table S2. Primers used in this study.**

| Primer | Sequence (5’-3’) |
|---|---|
| oTTR1 | TTGCCGCAGATTGAAGAG |
| oTTR2 | AGGTGGACACTGCAAATAC |
| oTTR3 | CGCGCTGACTCTGATATTATG |
| oTTR4 | CAAAGAGGAGCTGCTGTAAC |
